# Leveraging Transcriptomics Data for Genomic Prediction Models in Cassava

**DOI:** 10.1101/208181

**Authors:** Roberto Lozano, Dunia Pino del Carpio, Teddy Amuge, Ismail Siraj Kayondo, Alfred Ozimati Adebo, Morag Ferguson, Jean-Luc Jannink

**Affiliations:** School of Integrative Plant Science, Section of Plant Breeding and Genetics, Cornell University, Ithaca, NY; Department of Economic Development, Jobs, Transport and Recourses, AgriBio Centre for AgriBioscience, Bundoora, Australia; National Crop Resources Research Institute (NaCRRI), P.O. Box 7084, Kampala, Uganda; International Institute for Tropical Agriculture (IITA), Nairobi; United States Department of Agriculture, Agricultural Research Service (USDA-ARS) R.W. Holley Center for Agriculture and Health, Ithaca 14853, NY, USA

## Abstract

**Background:** Genomic prediction models were, in principle, developed to include all the available marker information; with this approach, these models have shown in various crops moderate to high predictive accuracies. Previous studies in cassava have demonstrated that, even with relatively small training populations and low-density GBS markers, prediction models are feasible for genomic selection. In the present study, we prioritized SNPs in close proximity to genome regions with biological importance for a given trait. We used a number of strategies to select variants that were then included in single and multiple kernel GBLUP models. Specifically, our sources of information were transcriptomics, GWAS, and immunity-related genes, with the ultimate goal to increase predictive accuracies for Cassava Brown Streak Disease (CBSD) severity.

**Results:** We used single and multi-kernel GBLUP models with markers imputed to whole genome sequence level to accommodate various sources of biological information; fitting more than one kinship matrix allowed for differential weighting of the individual marker relationships. We applied these GBLUP approaches to CBSD phenotypes (i.e., root infection and leaf severity three and six months after planting) in a Ugandan Breeding Population (n = 955). Three means of exploiting an established RNAseq experiment of CBSD-infected cassava plants were used. Compared to the biology-agnostic GBLUP model, the accuracy of the informed multi-kernel models increased the prediction accuracy only marginally (1.78% to 2.52%).

**Conclusions:** Our results show that markers imputed to whole genome sequence level do not provide enhanced prediction accuracies compared to using standard GBS marker data in cassava. The use of transcriptomics data and other sources of biological information resulted in prediction accuracies that were nominally superior to those obtained from traditional prediction models.

## Background

Genomic Selection (GS) [1] is a breeding method that exploits high-throughput genotyping technologies, novel statistical methods and the availability of genomic information. It has been used extensively in animal breeding and promises to impact plant breeding, particularly within clonally propagated and perennial plant systems [2].

GS approaches tend to avoid marker selection, and instead, all the marker information is utilized within the prediction models. Given such scenario where the number of predictors (*p*), is greater than the number of available observations (*n*) traditional regression models achieve poor predictive ability as a result of multicollinearity and overfitting among the predictors [2,3]. Several statistical methods have been explored to overcome these problems; shrinkage methods, where the regression coefficients are shrunk towards zero, are widely used for genomic predictions [4]. These methods include Genomic Best Linear Unbiased Predictions (GBLUP) [5], Bayesian regression [1,6], Least Absolute Shrinkage and Selection Operator (LASSO) [4] and ridge regression BLUP (rr-BLUP) [7]. Recently, machine learning methods have been proposed for genome-enabled predictions as they are capable of dealing with the dimensionality problem in a flexible manner [8,9]. Performance comparisons among these models have been conducted in several plant species [10–13] showing that the best statistical approach depends highly on the trait and the species that is being analyzed.

GS predictions rely on linkage disequilibrium (LD) between the markers and the Quantitative Trait Loci (QTL). Given the dramatic drop in sequencing costs, full-genome sequence data was proposed to be used in genomic predictions [14]. Simulation studies suggest that the use of whole genome sequence data would result in increased accuracy of genomic predictions [14–16] because the accuracy that can be achieved by the prediction model is no longer tied to the LD-QTL relationship as the causal mutations are present in the dataset [15].

Whole-genome sequencing is still prohibitively expensive for most crop breeding programs as the number of individuals evaluated can reach the tens of thousands. An efficient and cost-effective approach is to impute the whole-genome sequence variants of the individuals using a low-density genotyping platform and a previously sequenced reference population (reference panel) [17]. This system is widely used in human genetics, where large-scale sequencing efforts, like the 1000 Genome Project [18], provides standard reference panels for imputation.

In livestock and some crops, breeding populations are typically derived from a small group of common ancestors within a few generations in the past. Thus, these populations tend to have a small effective population size (Ne); this is a perfect scenario for performing whole genome imputation (WGI) as low-density markers will be able to adequately trace the haplotypes inherited from the ancestors [15] easing the imputation process.

Genomic prediction models tend to use unannotated anonymous markers, even when this is currently slowly changing, most models do not take into consideration whether SNPs are close to genic or regulatory regions. When imputing markers to whole sequence level, the number of predictors utilized increases significantly and so does the p >> n problem; this might prevent the model to put sufficient weight on the causal variants [19] thus affecting prediction accuracies. The use of biological priors has been proposed to both alleviate this problem and reduce the computational burden associated with models using millions of markers [20].

Over the last few years, several methods have been developed to incorporate biological or functional information into Association Studies and Genomic Prediction. In cattle, for example, Fortes et al. used an Associated Weighted Matrix (AWM) [21] to infer a set of genes related to beef tenderness. They later demonstrated that making genomic predictions with only SNPs near the inferred genes for beef tenderness resulted in prediction accuracies that were higher than when the entire marker set was used [22]. Other methods have sought to exploit biological information while avoiding marker selection. Su et al. [23] for example, tested a genomic BLUP (GBLUP) model where the relationship matrix was weighted using prior Bayesian models or GWAS summary statistics [23,24].

In contrast to the traditional GBLUP that assumes that all SNPs have the same effect-size distribution, methods like GFBLUP [25] or MultiBLUP [26] add one or multiple genomic random effects that quantify the importance of different marker sets respectively. These marker sets are typically defined by some source of biological evidence (i.e., metabolic pathway, sequence annotation, transcriptomics, evolutionary constraints).

A Bayesian method that has also been implemented to leverage biological information in prediction efforts is BayesRC [27] which uses a mixture of normal distributions to model SNP effects and include prior biological knowledge. BayesRC [28] allows the user to *a priori* allocate the SNPs into classes where each class is believed to have a different probability of containing causal variants for the trait. The aforementioned genomic feature modeling approaches (GFBLUP, MultiBLUP, and BayesRC) were designed to improve prediction accuracies of complex traits if the groups of markers selected are enriched for causal variants [28,29].

Transcriptomics studies have allowed researchers to investigate gene expression dynamics of different organisms in different tissues, conditions or developmental stages [30]. It can be of aid to discover genes and pathways that are involved in the regulation of complex traits, potentially revealing genomic regions that would be enriched in variants affecting specific traits [25,31]. Transcriptomics studies have already been used effectively as a source of biological priors to predict complex traits in cattle [20,25]. These studies showed that using informed models could slightly improve prediction accuracies when making same breed predictions and that the observed improvement was more evident with a greater genetic distance between the training and validation population (across-breed predictions).

Cassava (*Manihot esculenta*) is a major staple crop in parts of sub-Saharan Africa and is the primary source of calories for millions of people across the world [32]. Cassava Brown Streak Disease (CBSD) is a viral disease that hampers the production of cassava and is considered a serious threat to food security in Africa [33,34]. CBSD is caused by two distinct single-stranded RNA viruses, Cassava Brown Streak Virus (CBSV) and Ugandan Cassava Brown Streak Virus (UCBSV) [34–36]. Recently, transcriptomics data in cassava has been used to unravel the transcriptional dynamics of cassava plants under infection by both UCBSV [37] and CBSVs [38].

In the present study, CBSD phenotypes (root infection and leaf severity three and six months after planting) from a Ugandan Breeding Population (n=955) were analyzed using whole genome imputation (WGI) data (~million SNPs) and biological information coming from transcriptomics experiments [37,38], Genome-Wide Association Studies (GWAS) [39] and in-silico identification of immunity-related genes [40,41]. Our main objective was first to assess the feasibility of performing whole genome imputation in cassava and second to test if prediction accuracies can be enhanced by using WGI together with biological priors using GBLUP-derived models.

## Methods

### Plant material

Two diverse cassava populations were combined and used as a composite set for this study; individuals in this composite data set represented the genetic diversity of the Ugandan cassava gene pool. The first population (“Training“) was comprised of a panel of cassava accessions from the breeding program of the National Crops Resources Research Institute (NaCRRI) in Namulonge, Uganda. This population was the first used to train genomic prediction models for applied breeding at NaCRRI. The second population, (“GWAS“) was developed by Kayondo et al. [39] and was comprised of accessions. This population is derived from parents from the International Institute of Tropical Agriculture (IITA), The International Center for Tropical Agriculture (CIAT) in Colombia and some landraces of East Africa. Briefly, the “Training” panel was evaluated in two years (2012-2013), and three locations in an alpha-lattice design, and the “GWAS” panel was evaluated in a single year (2015) at three locations using an augmented randomized complete block design. For more information on both populations, please refer to [39]. For a list of the accessions used, see Table S1.

### Phenotyping Platform

The composite plant population was phenotyped for three separate traits: foliar CBSD severity measured three (CBSD3) and six (CBSD6) months after planting and CBSD severity in the storage roots (CBSDR) after a year. Briefly, CBSD severity was scored based on a 5-point scale with a score of implying an asymptomatic plant while a score of would mean over 50% of leaf vein clearing for foliar symptoms (CBSD3 and CBSD6) and 50% of root-core being covered by necrosis for CBSDR. Please refer to Kayondo et al. [39] for further details.

### Genotyping by sequencing and imputation

Genotyping-by-sequencing (GBS) libraries [42] were constructed as previously described [43]. Marker genotypes were called using the TASSEL 5.0 GBS discovery pipeline [44] after aligning the reads to the *Manihot esculenta* Version assembly. Genotype calls were stored in Variant Calling Format (VCF) files (one per cassava chromosome). The VCF files were filtered using VCFtools [45]; individual marker calls were masked if the read depth was lower than 3x, cassava genotypes with > 80% missing calls and SNP markers missing more than 60% were removed. Insertions, deletions, and multi-allelic markers were also withdrawn from the dataset. Beagle 4.1 software [46] with default parameter settings was used for imputation. In total 173k SNPs were called among individuals. This dataset was further filtered by an Estimated Allelic r-squared statistic (AR2) > 0.3 and a minimum Minor Allele Frequency (MAF) of 1%. The final set herein referred to as the “GBS” dataset, included 41,SNP markers called among the individuals.

### Imputation to whole-genome sequence data

Beagle 4.1[46] and Impute2[47,48] were tested and compared for imputation accuracy, marker density, and marker distribution. For both software’s, a Cassava Haplotype Map (HapMap) of accessions was used as a reference panel. This reference panel represented cultivated, hybrid and wild cassava relatives and contained million SNP markers [49].

#### Beagle Imputation

Imputation using Beagle 4.1 was performed in two steps (Figure S1). During the “BEAGLE Stage I” phase, a subset of the HapMap markers was used, including bi-allelic SNPs with MAF greater than 1%. Additionally, a 10bp thinning filter was set up, meaning that only one marker per 10bp was allowed. The resulting set included 716k markers with MAF > 1% and AR2 > 0.3. The BEAGLE Stage I marker set was then used in the second round of full HapMap imputation. The second marker dataset, “BEAGLE Stage II” had million markers exposed to the same MAF and ARfilters. The genetic positions of the HapMap markers were inferred using a smooth spline fit to the 22,403-marker composite map published by the International Cassava Genetic Map Consortium (ICGMC) [50]. The genetic positions were forced to be monotonically increasing, which is a requirement for BEAGLE to run properly. Beagle 4.1ran with default parameters. For this manuscript, only the ‘BEAGLE Stage II” markers were considered, and herein it will be referred to as the “BEAGLE” dataset.

#### Impute2 Imputation

Imputation using IMPUTE2 was performed in a single step (Figure S2). The number of haplotypes used as “custom” reference panel (−k_hap) was set to 400, the effective population size (Ne) to 1000, and the imputation window to 5Mb. The genetic positions of the HapMap were inferred as described in the “Beagle Imputation” section of this manuscript. The IMPUTE2 software, however, requires knowing the recombination rate between the current position and next position on the map. This recombination rate was calculated using the following formula:

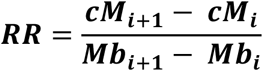

Where **cM** represents the genetic position of each marker “*i*” and **Mb** notes the physical position in megabases. The accuracy of the imputation was assessed using internally-calculated concordance tables. Briefly, IMPUTE2 masks the genotypes of one variant at a time from the study data (GBS markers) and then imputes the masked genotypes with information from the reference panel and the nearby variants. The percentage of concordance between the masked and the imputed genotypes for each 5Mb imputed window were subsequently calculated (Figure S3). Additionally, allele frequencies and imputation quality distributions were calculated and depicted by the IMPUTE2 information measure statistic “info” [48] (Figure S4) and imputation quality by allele frequencies (Figure S5).

### Biological Information

Three different sources of biological information related to CBSD resistance were used in this study.

#### Transcriptomics profiling

RNAseq data were obtained from two experiments. The first experiment [37] focused on profiling the transcriptome response across seven time-points after infection with UCBSV. Two contrasting cassava genotypes were used: ‘Namikonga’ (CBSD resistant) and ‘Albert’ (CBSD susceptible) (Figure S6). The libraries (Table S2) were checked for read quality using FastQC [51]. The Tuxedo Suite of programs [52,53] was then used to process the sequenced data. Reads in FASTQ formats were aligned to the *M. esculenta* reference genome v6[54] using TopHat v2.1.1/Bowtie v2.2.8[55]/[56]. A reference annotation of the cassava gene models (v6.1) from the Phytozome database was provided (https://phytozome.jgi.doe.gov). This version of the gene annotation contained a total of 33,033 transcripts. The minimum and maximum intron length were set to and 15,000bp respectively; the remaining parameters were set to default values. Subsequently, the *Cuffdiff* program within Cufflinks version 2.2.1 [57] was used to identify differentially expressed (DE) genes at each time-point among infected plants and controls. A false discovery rate of 0.01 after the Benjamini-Hochberg correction for multiple testing was used.

The second transcriptomics data was taken from Anjanappa et al. [38]. In this experiment, two cassava genotypes, the resistant ‘KBH 2006/18’ and the susceptible ‘60444’, were challenged against a mix of CBSV strains (CBSV - TAZ-DES-01 and UCBSV - TAZ-DES-02). RNAseq was performed days after infection; this time point was selected because it showed homogenous virus titer levels across the biological replicates in the susceptible genotype. Raw reads were not re-analyzed; a list of DE genes was extracted from the Anjanappa et al. manuscript (Table S3).

#### Quantitative Trait Loci

Kayondo et al. recently reported two major QTLs for CBSD foliage symptoms [39], one near the end of chromosome and another on chromosome that collocates with a previously reported, large introgression from wild cassava (Figure S7). Bi-parental QTL mapping has also identified hits on chromosomes and for foliar symptoms [58] and chromosome for root necrosis [59]. Small effect QTLs related to CBSD symptoms on roots were also detected, but they were not considered in this study.

#### Immunity-related genes

The most common disease resistance genes in plants are those belonging to the NBS-LRR family [60]. This highly conserved gene family has already been identified and positioned in a previous version of the cassava genome (Cassava Genome v5.0) [61]. In that study NBS-LRR and partial NBS-LRR genes were reported. Positions for each NBS-LRR genes were updated to fit in the latest cassava genome assembly (http://phytozome.gov, Cassava Genome v6) using Blast+ [62] (Table S4). Additionally, immune-related genes listed by Soto et al. [41] were added to this list (Table S4).

### Associating markers with genes

Markers that appeared within the coding region of a gene (defined as 5’UTR to 3’UTR, including introns) were considered to be “tagging” that gene. Bedtools [63,64] and in-house scripts (available from the GitHub page of this manuscript) were used to associate SNP markers to genes of interest.

### Co-expression Networks using WGCNA

Weighted Co-expression Network Analysis (WGCNA) [65,66] was used to identify highly correlated genes across different time-points based on their expression. Briefly, Fragments Per Kilobase of exon per Million reads (FPKMs) were log2 transformed. Genes without variation across the seven timepoints were filtered out using a Coefficient of variance (*CV = σ/µ*) cutoff of 0.9. Analyses were performed using the ‘WGCNA’ package in R programming software [67]. As previously described [66], ‘WGCNA’ calculates an expression Pearson’s correlation matrix for the genes, this matrix is later raised to a power β (0.8 in this study) before continuing with the clustering procedure. The ‘WGCNA’ *treecut* parameter was set to 0.85; the three parameters CV, β and *treecut* values were selected based on the number and quality of the co-expression modules identified. All other parameters were set to the package’s default values. To visualize the general trend of each module, eigengenes were calculated as the first principal component of the normalized expression values of all genes within a module and plotted as a heatmap [68,69].

### Genomic Selection Models

A two-step approach was used to evaluate genomic predictions in this study. This method was used to increase computational efficiency and control for differences in experimental design between different datasets. The first step involved accounting for trial-design variables using linear mixed models to calculate de-regressed Best Linear Unbiased Predictions (BLUPs), and the second step used the de-regressed BLUPs as phenotypes in the prediction model.

#### Genotypic value estimation

De-regressed BLUPs were calculated according to Garrick et al. [70]. The procedure has been described previously [12,71] and for this composite population specifically in Kayondo et al. [39]. Briefly, a mixed model was fit with the population mean and location as fixed effects and clone and breeding design variables (i.e., block, range) as random effects. BLUPs for clones represents an estimate of the total genetic value (estimated genetic value, EGV). Clone effect BLUPs (EGVs) were then extracted as the de-regressed BLUPs following:

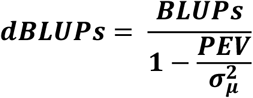

Where 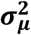 is the genetic variance and PEV is the prediction error variance of the BLUPs. Solutions for both 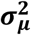 and PEV were retrieved from the mixed models solved using the *lmer* function of ‘lme4’ package [72] in R software.

#### Prediction models

We used three variations of the classic GBLUP to predict estimated breeding values (GEBV) for CBSD related traits:

**GBLUP** was fit using a linear mixed model of the form:

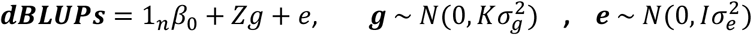

Where the solution for **g** represents the GEBVs. Briefly, ***β***_**0**_ is the mean, vector **g** is the random effect for the genetic markers, **Z** is a design matrix pointing observations to genotype identities, and ***e*** are the residuals. We assume that **g** has a known covariance structure defined by the genomic realized relationship matrix **K**. The genomic relationship matrix **K** was constructed using SNP dosages and an Rcpp [73] implementation of the function *A.mat* in the R package ‘rrBLUP’ [74]. GBLUP predictions ran using the function *emmreml* in the ‘EMMREML’ R package [75].

**GFBLUP** [29,76] is a modification of the traditional GBLUP that includes an additional genetic random effect; the linear mixed model followed the form:

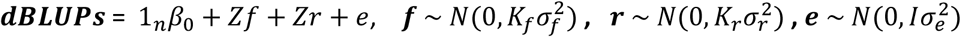

where ***K***_***f***_ and ***K***_***r***_ were genomic relationship matrices built using the SNPs within and outside the genomic feature. Specifically, ***K***_***f***_ was calculated with markers thought to be enriched for causal variants and ***K***_***r***_ was calculated with the rest of the markers in the genome. The relationships matrices were calculated as described before and the GFBLUP predictions were conducted using the *emmremlMultiKernel* function in the ‘EMMREML’ R package [75].

**MULTIBLUP** [26] was also used. This method is similar to GFBLUP but allows for multiple genetic random effects. As with GFBLUP method, predictions were conducted using the *emmremlMultikernel* function implemented in the ‘EMMREML’ R package.

#### Cross-validation

The accuracy of genomic prediction was measured as the correlation between the total genetic value (EGV, the random genetic effect from the first step regression model, not de-regressed) and the GEBVs. We used replications of a five-fold cross-validation scheme to obtain unbiased estimates of the prediction accuracies. The process of cross-validation used in this study was previously detailed by Wolfe et al. [13].

## Results

### Describing the population

We used the GBS marker dataset (~40K SNPs) to describe the LD patterns, population structure, and MAF distribution within a composite set of cassava varieties (Figure 1). After plotting the mean LD score (As in GCTA-LDS, [77]) of each variant, we noted a high level of LD heterogeneity across the entire cassava genome. Major LD peaks were not observed in centromeric regions, as would be expected with the common fall in recombination rate. Some high LD clusters were observed, however, near to the telomeres (Figure 1a). High LD across chromosome and at the end of chromosome were consistent with two relatively recent introgressions from a wild cassava relative [54]. The unique LD pattern in these two chromosomes was evident after plotting a regular LD decay plot (Figure 1b). Principal component analysis (PCA) on the dosage marker matrix (Figure 1c) indicated that there is little genetic differentiation between the two populations merged for composite analysis in this study. Moreover, the percentage of variance explained by the first two PCs was only 8.95%. The allele frequency distribution was also similar between the two populations (Figure 1d).

**Fig 1.**
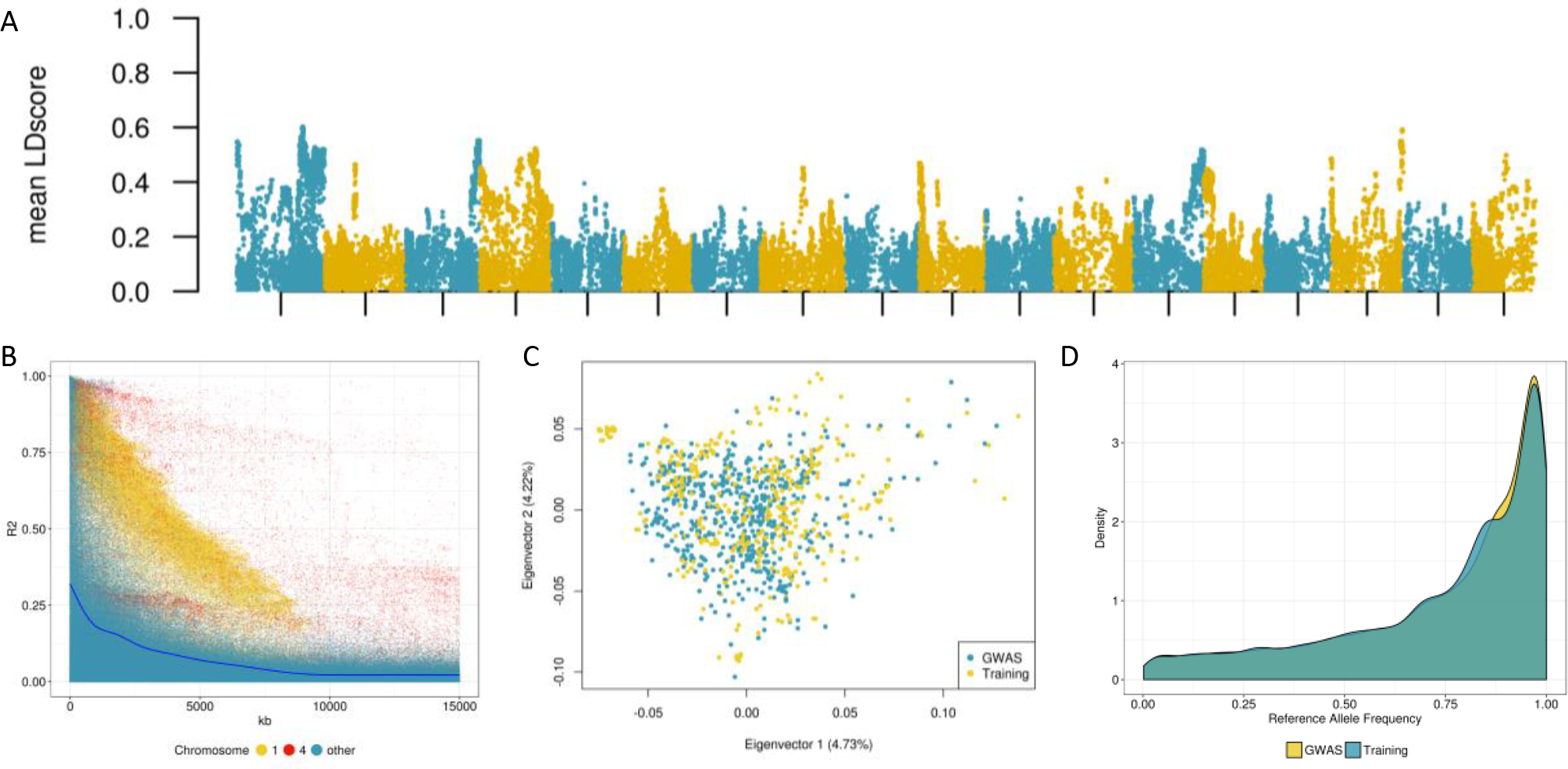
Describing the Breeding Population. **a** Local LD patterns across the cassava chromosomes as depicted by the mean LDscore of each marker. **b** LD decay plot, A random subset of all the r^2^ values of SNPs closer than 15Mb were plotted. Chromosomes and were plotted separately to highlight the distortion in their LD patterns due to the introgressions. **c** Principal component analysis using the SNP marker matrix, the two breeding populations that were merged in this study are shown in different colors. **d** Distribution of the reference allele frequencies between the two breeding populations.

### Imputation to whole genome sequence

We compared two different methods to impute the GBS dataset to a whole-genome sequence. BEAGLE and IMPUTE2methods have been challenged before regarding imputation accuracy and computational time, the results of which suggest that both approaches are sufficiently robust [78]. To select genetic markers that would “tag” candidate genes, we focused on the number and distribution of higher quality imputed SNPs (AR2/info > 0.3, MAF > 0.01) across the cassava genome. Using IMPUTE2 resulted in high-quality markers, more tagged genes (Figure 2a), and better marker distribution (Figure 2b, Figure S8) than BEAGLE. The total number of predicted genes in the current cassava assembly was 33,033. We tagged 32% of them using GBS markers, 70% using the BEAGLE imputed dataset, and 91% when using IMPUTE2. Other quality control tests were performed on the IMPUTE2 dataset, including imputation accuracies per chromosomal segments, distribution of allele frequencies, and “info” quality scores (Figure S3-S5).

**Fig 2.**
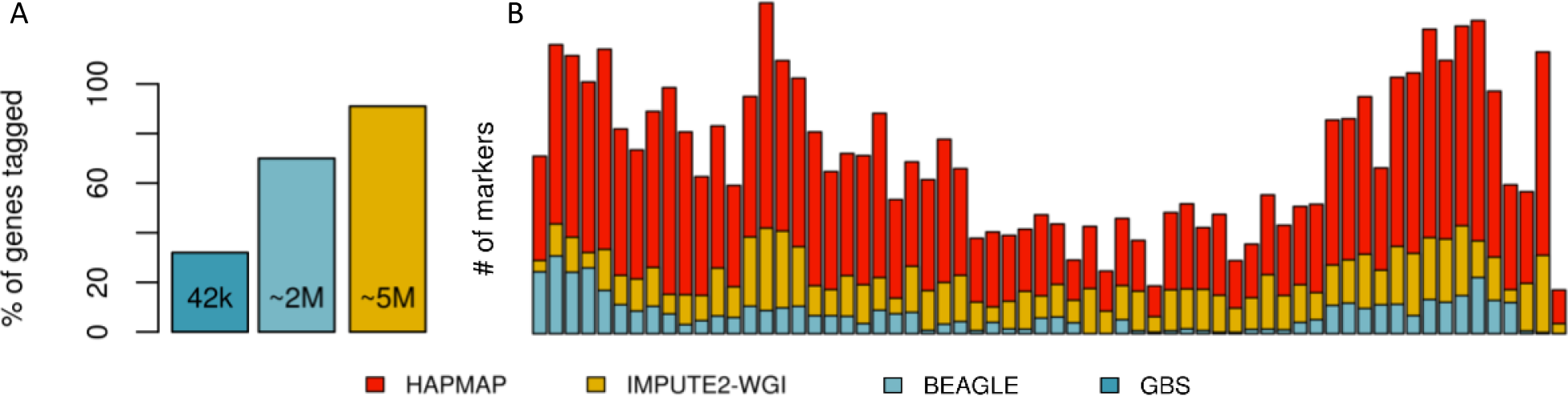
Imputation to whole-genome sequence. **a** Percentage of genes “tagged” using different SNP marker sets, the numbers inside the plots represents the number of markers. All markers considered had a MAF higher than 1% and an imputation quality value AR2/info higher than 0.3 **b** Marker distribution across chromosome 12, each bar represents a bin of 0.5Mb. The red colored bars represents the “true” distribution of variability as reported in the cassava HAPMAP, in orange, the distribution of the IMPUTEdataset (^~^5M markers) and in blue the Beagle dataset (^~^2M markers).

### Impact of Imputation level on Genomic Prediction accuracies

Prediction accuracies of a regular GBLUP model for three CBSD-related traits are shown in Figure 3. Specific conclusions regarding the impact of different imputation levels on prediction accuracies are not possible, as there is not a common trend among the three traits. We did note, however, that there was not a significant increase in prediction accuracy using different imputation levels. Moreover, when evaluating Cassava Brown Streak Disease severity six months after planting (CBSD6), the accuracy using only GBS data was consistently higher than any of the imputation methods tested. We also compared the prediction accuracies using one subset of markers from IMPUTE2 that matched the position of the GBS markers (Impute2GBS) and another subset using only SNPs imputed with the highest reliability (AR2/info > 0.9, Impute290, n = 371,524). Again, the prediction accuracies resulting from these subsets were nearly identical to those obtained using the full GBS and IMPUTE2 dataset (Figure 3).

**Fig 3.**
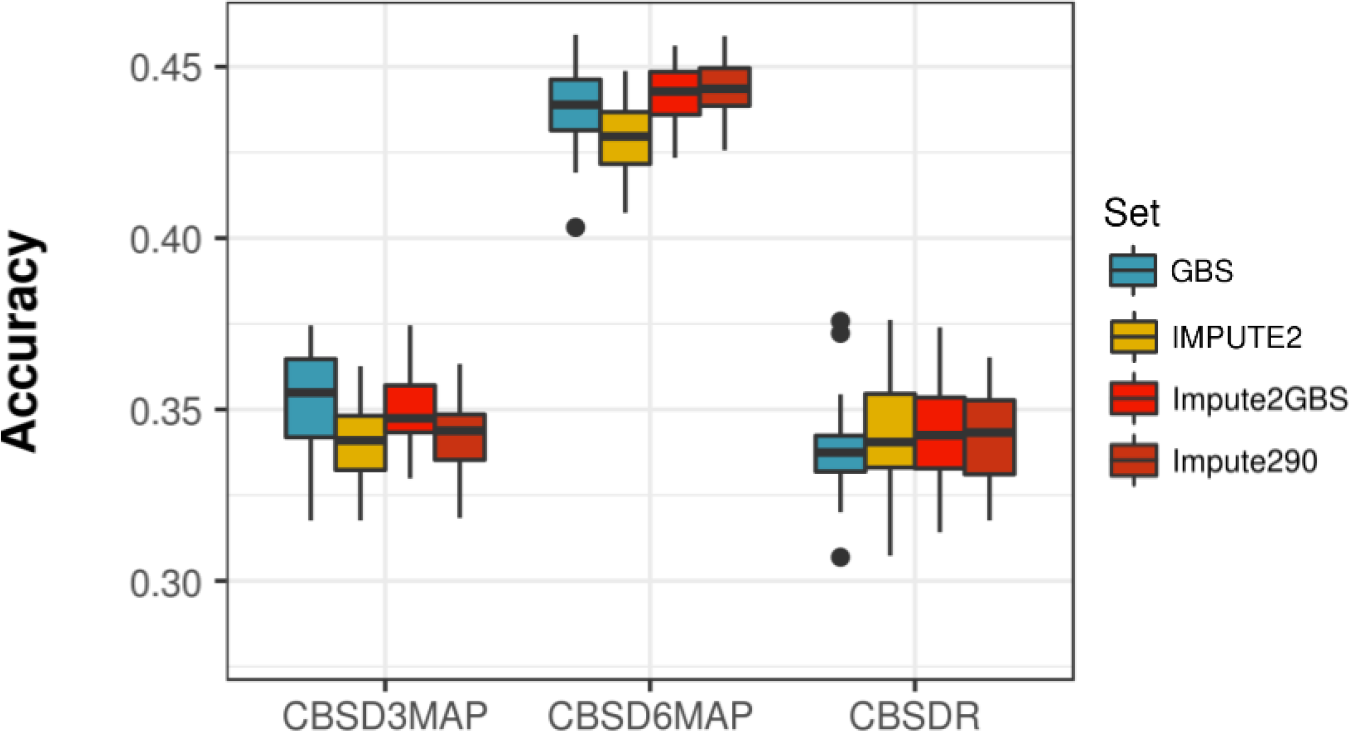
Impact of Imputation level on Genomic Prediction Accuracies. Comparing prediction accuracies for three traits; CBSD severity on leaves months after planting (CBSD3MAP), months after panting (CBSD6MAP) and CBSD severity on roots one year after planting (CBSDR) when using GBS (42k SNPs), the whole-genome sequence imputed datasets using IMPUTE(^~^5M) and also prediction accuracies for a subset of the IMPUTEmarkers matching the position of the GBS set (Impute2GBS) and only marker with an “info” imputation quality score higher than 0.9(Impute290)

### Accounting for known QTLs

Kayondo et al. [39] previously conducted a Genome Wide Association Study (GWAS) and identified two big effect QTLs for foliar CBSD severity using the same cassava population presented in this manuscript. The first identified QTL was very wide and located in the middle of chromosome 4. This QTL appeared to co-locate with a recent introgression from a wild cassava relative. The second QTL was located at the end of chromosome (Fig. S7).

This study sought to evaluate the relative importance of these QTLs for genomic prediction accuracy. We first ran a genomic prediction GBLUP model which included two genomic random effects: the first built with markers from chromosome and the second built with markers from chromosome 11. We compared the partial and total accuracies of this model with another two-kernel GBLUP model built with two random chromosomes, excluding chromosomes or (Figure 4). A clear difference in prediction accuracy was observed when chromosomes containing QTLs (blue) and random chromosomes (white) were compared. Since these QTLs were detected on foliar symptoms, we observed that the influence of chromosome and is higher in predictions of foliar phenotypes (CBSD6) than in necrosis on roots (CBSDR). Additionally, when we compared the total accuracy of the model including only the chromosomes with identified QTLs, we observed the prediction accuracy for CBSDS was very close to the model calculated using all cassava chromosomes. We then fit a model with three kernels (i.e. chromosome 4, and the rest of the genome) to investigate if there was any additional variance beyond the chromosomes containing the important QTL (Figure S9). The total prediction accuracy increased slightly for each measured trait, but it did not reach the accuracy level obtained when all markers were used in a single kernel model. This result suggests that marker partitioning is performed at the cost of prediction accuracy.

**Fig 4.**
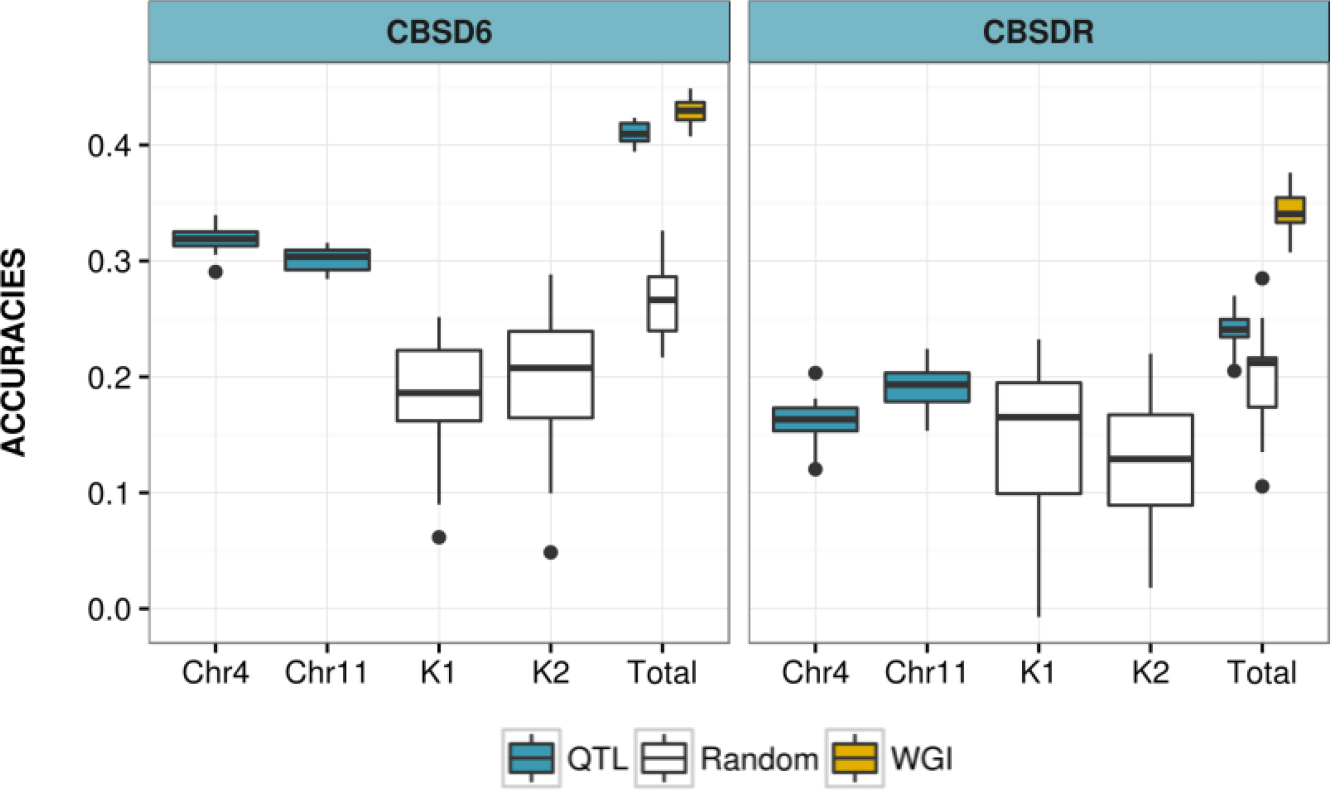
Accounting for the effect of previously reported QTLs. Comparing the maximum accuracy using whole-genome imputation (yellow) with two kernel GBLUP model using chromosome and only (blue) and random chromosomes excluding and (white) in each cross-validation iteration. Partial accuracies are shown under Chr, Chr11, Kand K2. Full model prediction accuracies are shown in “Total”. *CBSD6MAP: Foliar symptoms, CBSDR: Root symptoms.

### Using Transcriptomics data

Amuge et al. [37] profiled the response of two contrasting cassava genotypes to infection with UCBSV. RNA samples were collected across seven time points after inoculation by grafting with UCBSV and deep sequenced using the Illumina platform (Figure S6). Relative virus titer was quantified from the RNAseq libraries as the number of reads mapping to either CBSV or UCBSV genomes (Figure S10). Additionally, reads mapped to either of these genomes were de-novo assembled using Trinity [79] as a means of confirming the virus infecting the plant was only UCBSV and not CBSV (Figure S11). As previously demonstrated by Amuge et al., the transcriptional response of the two genotypes evaluated was radically different after UCBSV infection. While the tolerant cassava variety (‘Namikonga’) showed a strong response across most of the seven timepoints, the susceptible variety (‘Albert’) showed no transcriptional response between hours and days after infection (Figure 5, Table S5). Under the assumption that tagging and prioritizing SNPs close to genes contributing to the plant-virus interaction would increase prediction accuracies, we proceeded to explore different means of exploiting this dataset to locate these genes of interest.

**Fig 5.**
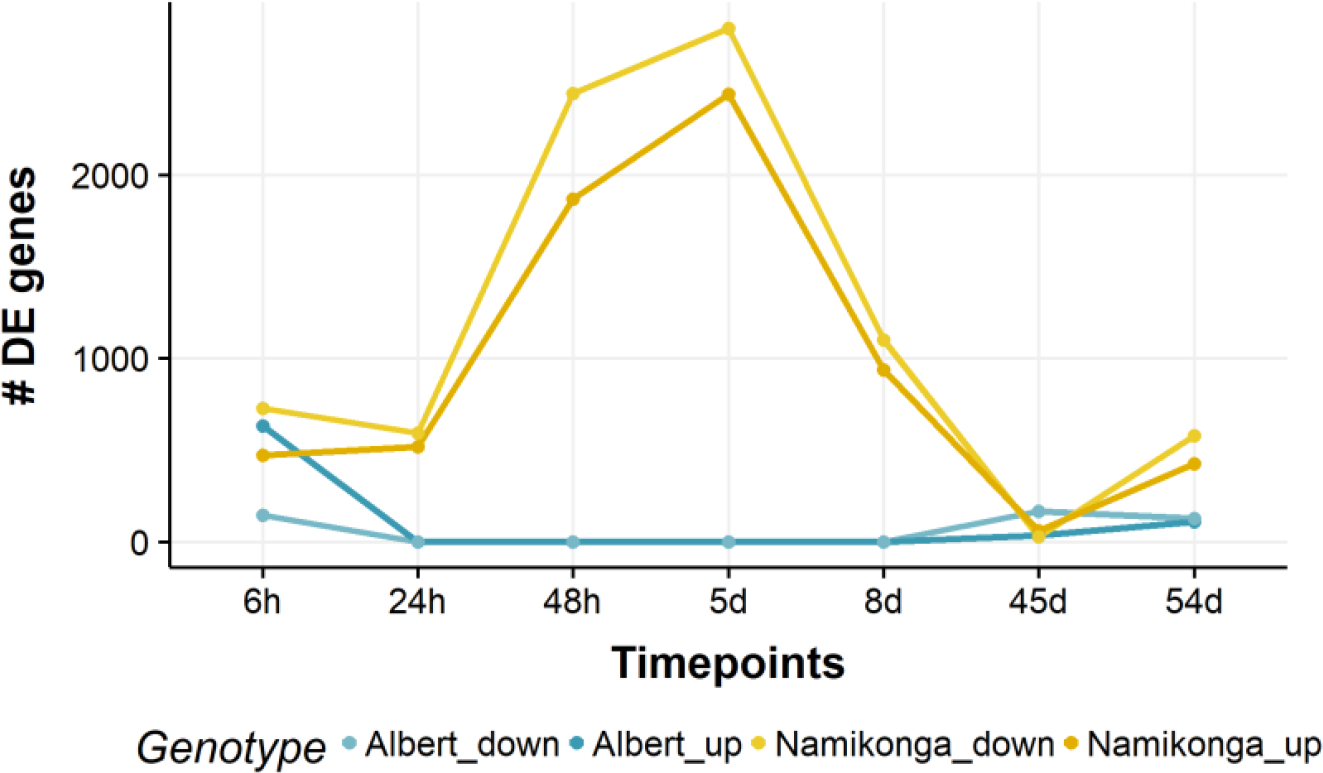
Transcriptional response to Infection with UCBSV. The test for differentially expressed genes was conducted at each timepoint between the infected an control plants using Cuffdiff. Genes considered to be differentially expressed had a q-value < 0.(Benjamini-Hochberg correction for multiple testing). *\*h = hours after infection, d = days after infection*

#### Differentially expressed genes

The most direct way to use the transcriptome dataset was to apply a GFBLUP procedure using the SNPs inside each Differentially Expressed (DE) gene as genomic features. We ran this analysis for two traits (CBSD6, CBSDR) and compared prediction accuracies between each GFBLUP model and the regular GBLUP model using the whole genome sequence imputed dataset (WGI) (Figure 6). In total, we ran eleven different GFBLUP models, including one comprised of DE genes across all time points (DE-all). While there were differences in the mean prediction accuracies between the models, none of them were significant.

**Fig 6.**
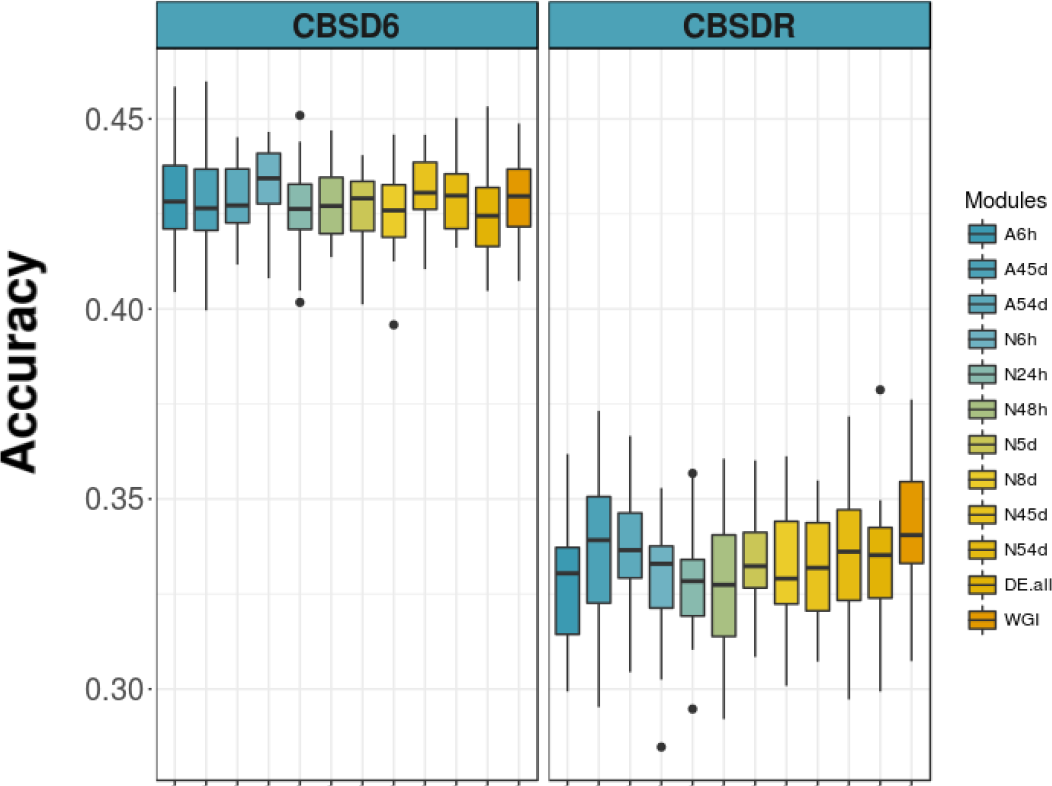
Using DE genes for Genomic Prediction. GFBLUP models (Two-kernel GBLUP) were fitted. For each model the Genomic feature kernel comprised SNPs inside the genes that were DE at each time point for each genotype. Three models for the susceptible DE genes (A6h, A45d and A54d), seven for the tolerant (N6h - N54d) and one for the combined DE genes (DE.all) were performed. Boxplot were the result of replications of 5-fold crossvalidation. \**A = Albert, N = Namikonga, h = hours after infection, d = days after infection*

#### Genes having a significant interaction between genotype and inoculation status

An alternative means of selecting genes of importance across all DE genes was to consider only those genes with a significant interaction with Genotype-by-Inoculation status (herein referred to as *GxI* genes). To accomplish this, a mixed model was fit for each gene:

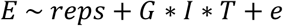

Where **E** is expression in FPKM, ***reps*** encompasses the three replicates as a random effect and ***G * I * T***describes the three-way, fixed effect interaction among inoculation status (***I***, infected or control), Genotype (***G***, susceptible or resistant) and the different time points (***T***). The p-values for each ***G * I*** interaction were extracted and corrected for multiple testing using a 5% FDR. Out of the total set of 33.033, genes in the cassava genome, 1,392 showed a significant *GxI* interaction at 5% FDR and at 1% FDR (Table S6). The genomic distribution of these genes appeared to be uniform (Figure 7a). When using GFBLUP, we noted that partitioning SNPs into two kernels based on whether they tagged *GxI* genes (at both 0.05 and 0.01 FDR thresholds) was not advantageous for prediction accuracies (Figure S12).

**Fig 7.**
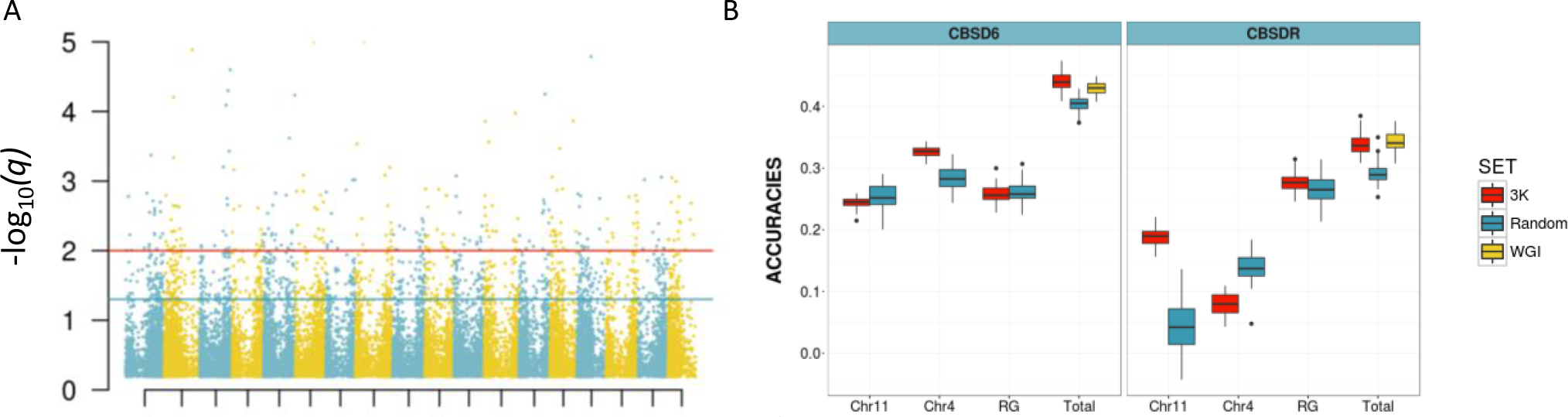
Filtering DE genes. **a** A linear mix model was used to calculate the genes that showed a significant interaction between inoculation status and genotype. In the manhattan plot −log_10_ “q-values” (FDR corrected p-values) for the *G x I* interaction term was plotted for the cassava chromosomes. The blue line is the threshold for 5% FDR and the red one for 1% FDR **b** Genomic predictions using three kernel GBLUP models. In red, the partial prediction accuracies (Chr11, Chrand RG) and total accuracy using only markers associated with significant GxI genes are compared with a three kernel model of random SNPs in blue and the regular single kernel GBLUP prediction using all the markers in yellow.

Based on previous results demonstrating the importance of large-effect QTLs on chromosomes and 11, we partitioned the *GxI* SNPs into three kernels: chromosome 4, chromosome and the rest of the genome. In this model, only SNPs inside the significant *GxI* genes (5% FDR) were considered. This was in contrast to the GFBLUP approach, where a kernel with information from the rest of the genome was fit. Thus, the number of SNPs used was much lower than the GFBLUP approach. The prediction accuracies using this three-kernel model were similar to those using the WGI dataset, despite using less than 2% of the SNPs (Figure 7b). To test that the *GxI* associated SNPs were relevant for prediction, we also ran a model using a different random set of SNPs during each of each of the rounds of cross-validation. These random SNPs were in approximately linkage equilibrium with the GxI-associated SNPs. The GxI-associated SNPs showed significantly better prediction accuracies than when random SNPs were used (Figure 7b). Given the apparently good results using the three-kernel method, we fit the same model with an extra kernel to account for the rest of the genome and while we expected an additional boost in prediction accuracies, we did not observe an increase (Figure S13). Whether the rest of the genome SNPs has spurious associations that decrease prediction accuracies or if there is an implicit “cost” for partitioning the genome in a multiBLUP model, are hypotheses that were not tested in this manuscript.

#### Co-expression modules

We used Weighted Gene Correlation Network Analysis (WGCNA) [65,66] to identify correlated genes based on their expression patterns across the different timepoints. WGCNA allows the identification of modules of genes that are more correlated within each other than they are to genes outside the module [65]. This unsupervised method was used to identify modules of co-expressed genes and test if any of these modules were more important or enriched in causal variants, the result of which would increase prediction accuracies for any of the CBSD related traits under a GFBLUP framework.

Of the 33,033 total genes in the reference cassava genome, 5,574 passed an ad-hoc Coefficient of Variance filter (*CV* = 0.9) and were used in downstream analysis. From the remaining 5,574 genes, 2,789 were assigned to modules containing between and genes (Table S7). A total of 2,785 genes could not be assigned to any module (Grey module). Eigengenes for each module were calculated and plotted in a heatmap depicting modules as rows and the timepoints, genotypes, and inoculation status as columns (Figure 7a). While some modules are noisy with a broad co-expression pattern across different timepoints and conditions, some of them are correlated at only one or two conditions (yellow, etan, and green). Other modules are dependent on time after infection, regardless of genotype or inoculation status (turquoise). Interestingly, two modules (black and cyan) grouped genes with ‘Namikonga’ and ‘Albert’ specific expression across all timepoints (Figure 7a).

We then used the identified modules to fit a GFBLUP model for each module. The accuracies obtained are shown in Figure 7b. For CBSD severity six months after planting (CBSD6) and severity on roots (CBSDR), none of the GFBLUP models provided a significant advantage in prediction accuracy over the traditional GBLUP (WGI). For CBSD severity three months after planting (CBSD3), however, one of GFBLUP module model (red, genes, 3,558 SNPs) obtained a prediction accuracy higher than WGI. Using WGCNA as a proxy to identify genomic features helped to marginally improve the genomic prediction accuracy for only one of the traits tested.

#### Other biological data

As a final step in this analysis, we incorporated all the available biological information, including large-effect QTL peaks, *GxI* genes, and previously identified immunity-related genes. The immunity-related genes included NBS-LRR genes[40], immunity-related genes as annotated by Soto et al. [41], and DE genes proposed to have a major role in the resistance response against joint UCBSV and CBSV infection in a single-point transcriptomics study (Table S3) [38].

Multi-kernel GBLUP models were fit with SNPs tagging each biological information category; chromosome large-effect QTL, chromosome large-effect QTL, *GxI* significant genes, and immunity related genes (Fig 8). A small increase in prediction accuracy for each of the traits was obtained through various combinations of the information above. For CBSD3, a three-kernel model with the chromosome large-effect QTL, tagged *GxI* genes, and genes present in the red WGCNA module increased accuracy by 1.7% (Fig 8a). For CBSD6, a four-kernel model using QTLs from both chromosome and chromosome 4, tagged *GxI* genes, and the immunity-related genes resulted in a 2.52% increase in prediction accuracy (Fig 8b). Finally, a three-kernel model considering only the chromosome large-effect QTL, the immunity related genes, and the tagged *GxI* genes resulted in a prediction accuracy increase of 2.52% for roots phenotyped one year after planting (Fig 8c).

**Fig 8.**
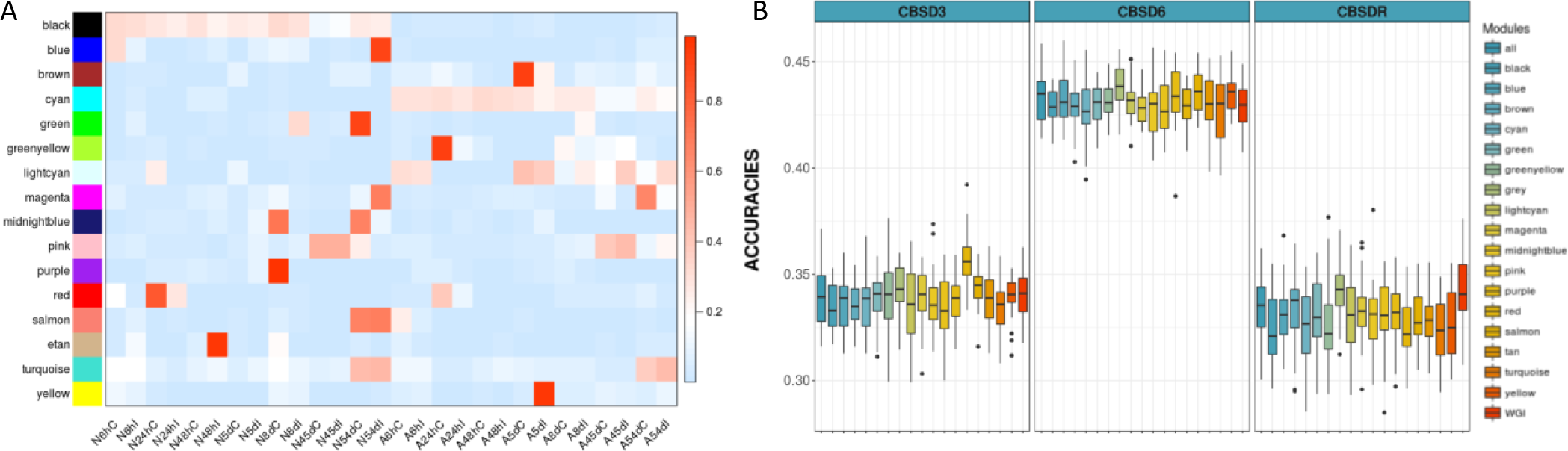
Co-expression network analysis. **a** Heatmap of eigengenes representing each co-expression module as obtained by WGCNA. All timepoints for both genotypes including controls were included and presented as columns. The identified co-expression modules are presented in each row. The eigengene values are a relative measure of expression levels of the genes in the module. **b** GFBLUP predictions using the modules information. As in figure the genes in each module were used to build a GFBLUP model, one kernel using SNPs within each module genes and the other covering the rest of the genome. Total prediction accuracies were plotted.

**Fig 9.**
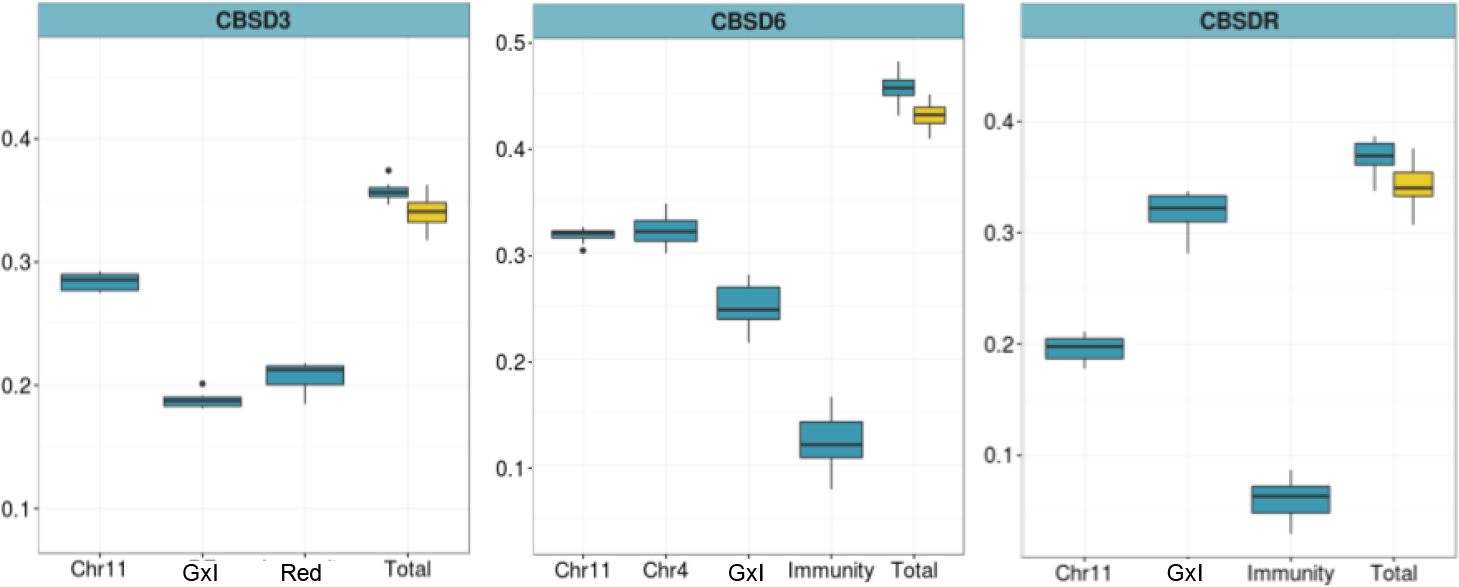
Combining sources of evidence. Four and three kernel GBLUP model including markers surrounding previously reported QTLs (chrand chr11), GxI genes found in this study, immunity related genes and the red WGCNA module (blue). Partial and Total accuracies are compared with the regular GBLUP model (yellow). A nominal increase in prediction accuracy of 1.7%, 2.52% and 2.5% was found for CBSD3, CBSDand CBSDR respectively.

## Discussion

In this study, we explored the improvement of genomic prediction in cassava through the integration of transcriptomics data, the genetic architecture of CBSD, biological priors, and whole sequence variants. Our results provide insight on how incorporating biological information into prediction models can impact genomic prediction within this important staple crop. Also, we explored models which can be extended to its use on other sources of biological data such as regulatory elements, evolutionary conserved regions, chromatin accessibility assays, and eQTLs.

### SNP imputation to Whole-genome sequence

Compared to the prediction accuracies obtained using GBS markers, imputed sequence data produced no advantage when applied to CBSD related traits. This behavior has been noted in other animal empirical studies, where marginal [80] or absent increases in prediction accuracy and reliability were observed [19,81–83]. Simulation studies, however, have reported significant gains in prediction accuracy under some circumstances (i.e., low MAF of the causal variants) [14–16]. As reviewed before [19], several reasons may account for this lack of increase in prediction accuracy when using imputed sequence data. Problems with the imputation method itself, small reference panels, and causal variants with low MAF may result in difficulties imputing sequence data. Additionally, many markers could result in models failing to put sufficient weight on the causal variants (i.e. a severe “*p* >> *>n*” problem).

In our study, an imputation reference panel of only 240 individuals was used to impute a dataset of 955 highly related individuals from NACRRI (Namulonge, Uganda). Additionally, the cassava genome has at least two major and recent introgressions from wild relatives [54] on chromosomes 1 and 4. Since wild cassava individuals are underrepresented in the reference panel [49] introgressed regions showed a significant drop in imputation accuracies (Fig S3). Moreover, the overall imputation accuracy in this dataset was significantly lower than when a larger and more diverse target panel was used. While these factors have affected the prediction accuracies, the purpose of using imputed sequence data in this study was to tag the maximum number of genes rather than just increase predictive accuracies by imputing to sequence level. That is, imputation was performed as a means of ensuring relevant genes could be tagged and used as additional information in the model.

### Genetic Architecture of CBSD

Genetic architecture of a trait is an important consideration when implementing different genomic prediction models. Genetic architecture can vary drastically from trait to trait but also from species to species. For example, in maize, most agronomic traits are controlled by many small effect loci. This is in contrast to rice, where many agronomic traits, including grain yield, have large effect QTLs [84].

Resistance to CBSD in cassava was historically considered to be a quantitative trait under the control of several contributing loci. However, large-effect QTLs were recently detected using association studies in a diverse population [39] and by traditional bi-parental QTL mapping [58,59]. In the present study, we showed that when genomic predictions were performed using only markers belonging to chromosomes containing the large-effect QTLs (i.e. chromosomes and 11), nearly the same prediction accuracies were obtained as when markers across the genome were used (Fig 4a). Since these QTLs were originally detected in leaves, it was no surprise that the prediction accuracies were not as high when the same models were used to predict CBSD severity on roots (Fig 4b). These data suggest an absence of correlation between root and shoot symptoms in cassava plants affected by CBSD. This phenomenon has been previously described; infected plants may show severe shoot symptoms and mild root necrosis or vice versa [85]. Moreover, the severity of symptoms has been demonstrated not to be correlated with virus titer, especially for resistant or tolerant varieties [85].

Previous research has tackled the problem of incorporating genotype-phenotype associations to boost genomic prediction by either adding significant markers as fixed effects [86,87] or by weighting the Genomic Relationship Matrix (GRM) with marker association information [88,89]. While we did not focus on any of these methods, tracking known QTLs allowed us to utilize better the information obtained from the transcriptomics experiment.

### On using Transcriptomics to Aid Genomic Prediction

Transcriptomics data has been used before as a source of biological priors for genomic prediction in cattle [25,28]. Like in the present study, Fang et al. [25] used transcriptomic regions responsive to Intra Mammary Infection (IMI) to fit a GFBLUP model that included a separate genomic effect of SNPs within DE genes. Similarly, MacLeod et al. used a novel Bayesian method (BayesRC), that allowed the incorporation of biological information by defining classes of variants likely to be enriched for causal mutations [28]. Both studies showed a minimal increase in prediction accuracies for within-breed predictions and a true benefit was observed only with across-breed predictions.

In this study, we analyzed existing transcriptomic data using three different approaches to explore multiple hypotheses related to the introgression of transcriptomics into genomic prediction models. The first approach exploited DE genes specific to each measured disease timepoint and cassava genotype (i.e DE genes six hours after infection in Namikonga) to fit a series of GFBLUP models. This approach explored whether any timepoint-genotype combination would be more enriched for causal variants and, thus, more useful for improving prediction accuracies. No increase in prediction accuracy was observed. This result was expected as we did not expect the response of individual genotypes to be representative of the entire population. Further, there were a total of 9,379 DE genes found in at least at one time point; this is close to one-third of the entire predicted gene set in the cassava reference genome.

To narrow the number of DE genes, we then hypothesized that genes exhibiting a significant statistical interaction between inoculation status (Control vs. Infected) and genotype (‘Namikonga’ vs. ‘Albert’) might be more relevant for CBSD related traits. Only 1,391 genes were significant to *GxI* (q < 0.05), and, while the multi-kernel GBLUP models performed better than when selecting the same number of random genes, the prediction accuracy remained the same as the full GBLUP model.

Finally, we used WGCNA to infer modules of co-expressed genes within the RNAseq dataset. This method has been used in several organisms to identify biologically meaningful gene modules, and it has helped to generate useful insights into how genes interact under certain conditions [66,69,90–92]. We assumed that modules consisting of highly interconnected genes would be enriched in causal variants and promote an increase in prediction accuracy under a GFBLUP framework. Only one module for one trait (red, CBSD3), however, showed a marginal increase in prediction accuracy

There are many reasons why we think the approaches using transcriptomics did not result in larger increases in prediction accuracy. First, the RNAseq data came from only two cassava varieties, and its transcriptome response may not be representative of the composite set used in this study. Secondly, samples were collected during the early (i.e., <54days) response of the plant to the infection. In contrast, the phenotypes were collected in the field three, six, and twelve months after planting. Thirdly, the plants were infected with only UCBSV (as confirmed by de-novo assembly of the viral reads, Fig S11), while under field conditions it is common to observe co-infection of CBSV and UCBSV [93]. Anjanappa et al. [38] previously showed that the response of cassava to a combined CBSV and UCBSV infection was significantly stronger in the susceptible variety than in the resistant variety. These results are in contrast to the current study, where ‘Namikonga’ showed a stronger response when only infected by UCBSV. As such, we can infer that the transcription response of cassava plants infected only with UCBSV may not be representative of infected plants in the field. Fourth, Increasing the accuracy of predictions using closely related individuals with long-range LD might not be an easy task in future breeding efforts. Rather, genomic prediction methods that incorporate biological priors may be more beneficial in across-breed prediction, where the LD structure is disrupted [28,76,82]. Specifically, Fang et al. found only a small increase (3.2% to 3.9%) in prediction accuracies by using GFBLUP and transcriptomics data when predicting milk traits within Holstein cows; the same study observed a 164% gain in prediction accuracy when the prediction was performed across-breeds.

Cassava Brown Streak Disease is currently present only in East and Southern Africa. Thus the Western African material cannot be evaluated for resistance to this disease because of the dangers of propagating the disease. In this scenario, a genomic selection model might be trained in the eastern African population(s) to predict resistance to CBSD in western germplasm. While these populations are not as divergent as cattle breeds, we expect that the LD structure between these two populations would be weaker and thus favor a model that uses prior biological information.

## Conclusions

The Genomic Prediction approach using prior biological information and markers imputed to whole-genome sequence achieved only a marginal increase in the accuracy of prediction for CBSD related traits. We believe that additional functional genomics research together with bigger reference panels that would improve imputation accuracies and a more precise phenotyping platform are necessary to unlock the potential of biology-assisted prediction models. Moreover, we think that this kind of novel approaches would provide insights into the genetic mechanisms underlying quantitative traits.

## Abbreviations

GS: Genomic Selection
GWAS: Genome-Wide Association Studies
CBSD: Cassava Brown Streak Disease
CBSV: Cassava Brown Streak Virus
UCBSV: Ugandan Cassava Brown Streak Virus
GBS: Genotyping-By-Sequencing
BLUP: Best Linear Unbiased Prediction
GEBV: Genomic Estimated Breeding Values
LD: Linkage Disequilibrium
SNP: Single Nucleotide Polymorphism
DE: Differentially Expressed
EGV: Estimated Genetic Value

## Declarations

### Ethics approval and consent to participate

Not applicable.

### Consent for publication

Not applicable.

### Availability of data and material

Sequences for every gene presented in this article are available in the Phytozome v10.repository, http://phytozome.jgi.doe.gov (*Manihot esculenta* v6.1). Scripts used in this manuscript are available at, https://github.com/tc-mustang/CBSDTrancriptomics. Dosage matrices and Variant Call Format (VCF) files can be accessed upon request through a secure FTP server. The transcriptome data from Amuge et al. [37] is available in the SRA BioProject ID PRJNA360340.

## Competing Interests

The authors declare that they have no competing interests.

## Authors’ contributions

The study was conceived and designed by RL, DPC and JLJ. AO and IK were in charged of data collection. MF and TA advised on CBSD and performed the transcriptomics study. The data analysis was performed by RL. The manuscript was written by RL and DPC. JLJ critically revised the manuscript with important scientific and statistical content All authors read and approved the final manuscript.

## Funding

This work was supported by the project “Next Generation Cassava Breeding Project” through funds from the Bill and Melinda Gates Foundation and the Department for International Development of the United Kingdom.

## Acknowledgements

To the memory of Martha Hamblin, whose friendship, guidance, and patience were paramount to this work. We would like to thank Deniz Akdemir for Statistical consulting.

